# eRNA-IDO: a one-stop platform for identification, interactome discovery and functional annotation of enhancer RNAs

**DOI:** 10.1101/2023.12.19.572028

**Authors:** Yuwei Zhang, Lihai Gong, Ruofan Ding, Wenyan Chen, Hao Rong, Yanguo Li, Fawziya Shameem, Korakkandan Arshad Ali, Lei Li, Qi Liao

## Abstract

Increasing evidence proves the transcription of enhancer RNA (eRNA) and its important role in gene regulation. However, we are only at the infancy stage of understanding eRNA interactions with other biomolecules and the corresponding functionality. To accelerate eRNA mechanistic study, we present the first integrative computational platform for human <u>eRNA</u> identification, interactome discovery, and functional annotation, termed eRNA-IDO. eRNA-IDO comprises two modules: eRNA-ID and eRNA-Anno. Functionally, eRNA-ID identifies eRNAs from *de novo* assembled transcriptomes. The bright spot of eRNA-ID is indeed the inclusion of 8 kinds of enhancer makers, whose combination enables users to personalize enhancer regions flexibly and conveniently. In addition, eRNA-Anno provides cell/tissue specific functional annotation for any novel and known eRNAs through discovering eRNA interactome from the prebuilt or user-defined eRNA-coding gene networks. The pre-built networks include GTEx-based normal co-expression networks, TCGA-based cancer co-expression networks, and omics-based eRNA-centric regulatory networks. Our eRNA-IDO carries sufficient practicability and significance for understanding the biogenesis and functions of eRNAs. The eRNA-IDO server is freely available at http://bioinfo.szbl.ac.cn/eRNA_IDO/.

## Introduction

The past decade has seen increasing evidence confirming the pervasive transcription of noncoding RNA from active enhancer regions, termed enhancer RNA (eRNA). Due to the dynamic nature of enhancer activity across different tissues and lineages, eRNA transcription shows high specificity to biological contexts [1]. eRNAs, once regarded as “transcription noise” or “byproduct” [2], have now been widely validated to play important roles in diversified biological functions and diseases such as cardiovascular development [3] and cancer [4]. Mechanistically, eRNA can promote enhancer-promoter loops and regulate epigenetics through interacting with several general factors, including components of cohesion or mediator [5, 6], and histone acetyltransferases CBP/p300 [4, 7]. In addition, the interaction of eRNAs with transcription elongation factors can facilitate the pause-release of RNA polymerase II pause-release to control transcription elongation.

Due to the increasing attention to eRNA functionality, several databases have been designed to characterize the transcription and potential targets of eRNAs, such as HeRA [8], TCeA [9], Animal-eRNAdb [10], and eRic [11]. However, these databases only provided information on annotated eRNA loci and enhancer regions, where users cannot investigate novel eRNAs. Furthermore, despite the existence of many ncRNA functional annotation platforms, they are not well-suited to eRNAs. For example, ncFANs v2.0 [12] requires known ncRNA identifiers as input, but in fact eRNAs have no reference ID or symbol. AnnoLnc2 [13] predicts the functions of novel lncRNAs based on co-expression networks. Still, neither considers cell/tissue specificity nor provides the eRNA-specific characteristics such as histone modification, chromatin architecture, and interactive molecules. Until now, a comprehensive platform for eRNA functional annotation is still lacking.

Therefore, we present the first one-stop platform for human <u>eRNA id</u>entification, interactome discovery, and functional ann<u>o</u>tation, termed eRNA-IDO (**Figure 1**). eRNA-IDO comprises two available modules: eRNA-ID and eRNA-Anno. eRNA-ID enables users to define enhancers and identifies enhancer-derived noncoding RNAs from the uploaded *de novo* assembled transcriptome. eRNA-Anno predicts eRNA functions by discovering eRNA-connected protein-coding genes (PCGs) in normal/cancer co-expression and eRNA-centric regulatory networks. All functions of eRNA-IDO can be realized based on pre-built data and also allow for user-defined data, thus carrying sufficient practicability and convenience for biological researchers. This web server is freely available at http://bioinfo.szbl.ac.cn/eRNA_IDO/ and opens to all users, without a login requirement.

**Figure 1.**
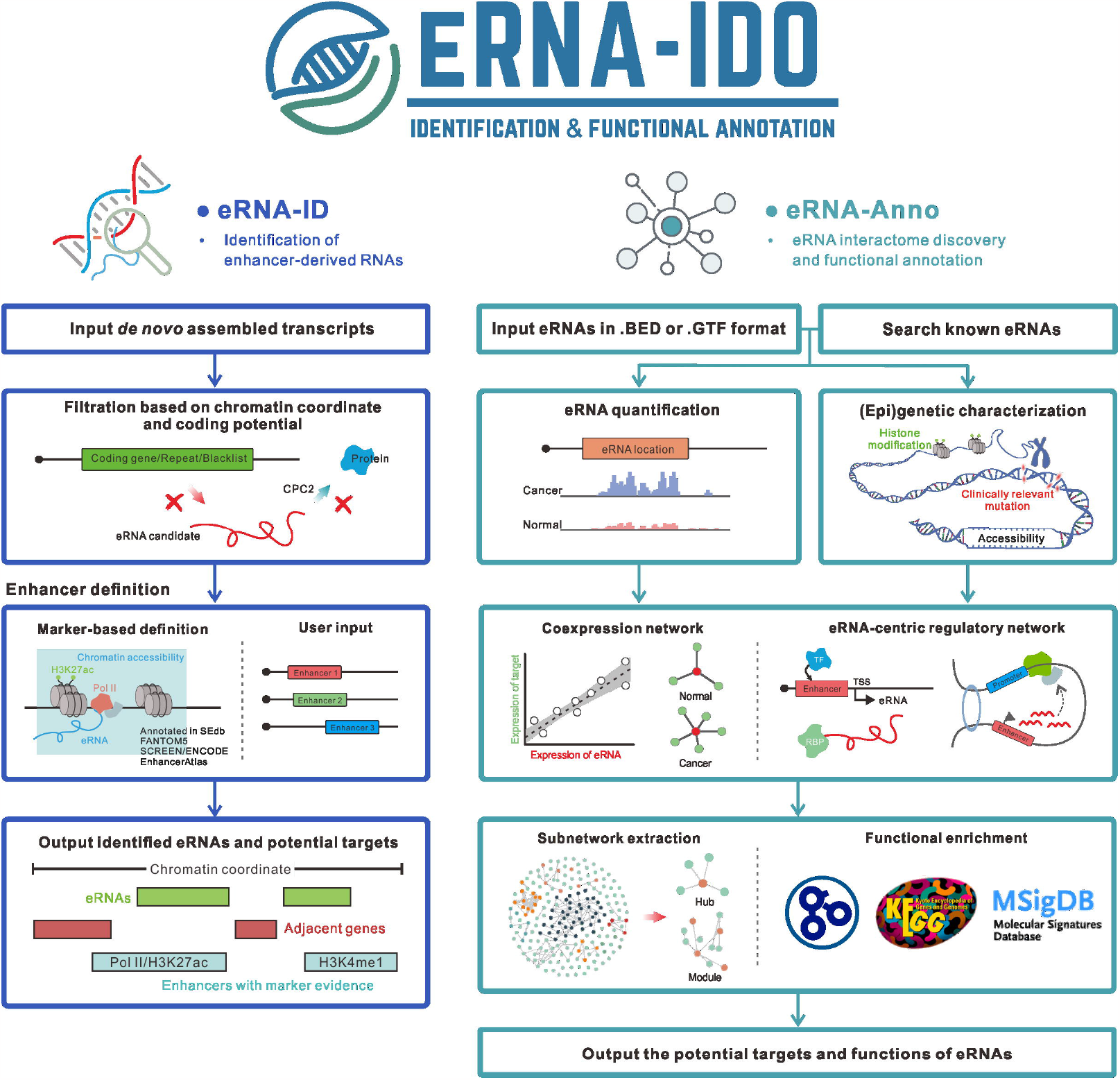
The workflow of eRNA-IDO. eRNA-IDO comprises two functional modules: eRNA-ID for eRNA identification and eRNA-Anno for interactome discovery and functional annotation.

## Materials and Methods

### Workflow and data architecture of eRNA-ID

The left panel of **Figure 1** shows the schematic workflow of eRNA-ID. eRNA-ID takes *de novo* assembled transcripts of RNA-seq or GRO-seq data provided by users as input. The transcripts overlapped with annotated PCGs, simple repeats, and blacklisted regions are removed according to the GENCODE v33 reference [14]. Next, the coding potential of the remaining transcripts is evaluated by CPC2 [15] (default parameter), and noncoding RNAs transcribed from enhancer regions are predicted as eRNAs. Enhancer regions can be either uploaded by users in BED format or defined using our marker buffet. The marker buffet is composed of 8 kinds of enhancer markers, including H3K27ac (**Supplementary Table S1**), H3K4me1 (**Supplementary Table S2**), chromatin accessibility (**Supplementary Table S3**), RNA polymerase II binding (**Supplementary Table S4**), super-enhancers from SEdb v2.0 [16], EnhancerAtlas v2.0 [17] enhancers, FANTOM5 [18] enhancers and SCREEN/ENCODE [19] enhancers, which are optionally overlapped or merged (using bedtools multiinter/merge) to obtain high-confidence or comprehensive enhancer profiles. The +/-3□kb regions around the center of the selected markers are defined as potential enhancer regions. These markers are cell/tissue-specific except those from FANTOM5 and SCREEN database. The data type, source, and number of biosamples of these enhancer markers are listed in **Table 1**. Finally, eRNA-ID outputs the chromatin locations, adjacent genes (+/- 1Mb), and enhancers of predicted eRNAs.

### Workflow and data architecture of eRNA-Anno

The right panel of **Figure 1** shows the schematic workflow of eRNA-Anno. eRNA-Anno takes either chromatin coordinates of novel eRNAs or the identifiers of known eRNAs annotated in HeRA [8] and eRic [11] database as input. Novel eRNAs should be input in BED or GTF format. For known eRNAs, the ENSR identifiers, chromatin coordinates, and adjacent genes (within +/-1Mb) are acceptable. Below is the detailed description for each procedure.

#### 1. eRNA quantification

The expression levels of known eRNAs are obtained from HeRA and eRic. Suppose chromatin coordinates of novel eRNAs serve as input, in that case, RNA-seq data from TCGA (https://portal.gdc.cancer.gov/) and GTEx portal [20] are used to quantify eRNA expression. To save the processing time, the expression levels are estimated based on the read coverage from BigWig files. The formula is:

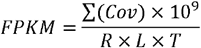

Where ∑(*Cov*) is the total read coverage of a given eRNA region, *R* is read length, *L* is eRNA length, and *T* is the total mapped reads of the library.

#### 2. Profiling genetic/epigenetic landscape

eRNA-Anno portrays a genetic/epigenetic landscape for eRNAs, including chromatin accessibility, clinically relevant mutation, and histone modification (H3K27ac and H3K4me1). Histone modification and chromatin accessibility are characterized based on ChIP-seq and ATAC-seq/DNase-seq from Cistrome database [21] (**Supplemental Table 1-3**). Clinically relevant mutations within the query eRNA regions are collected from ClinVar [22] and COSMIC [23] database.

#### 3. eRNA-PCG network construction

eRNA-Anno constructs a co-expression network between eRNAs and PCGs and an eRNA-centric regulatory network. Both user-uploaded expression matrix and publicly available data are supported for the co-expression network. Publicly available data refer to RNA-seq data of 52 normal tissues from GTEx portal [20] and 31 cancer types from the TCGA portal (**Supplementary Table S5**). The toolkit GCEN [24] calculates Spearman correlation coefficients and adjusted p-values.

For the eRNA-centric regulatory network, the relationships of eRNAs with TF, RBP, and E-P loop are investigated. eRNA-TF interactions are obtained based on 11356 ChIP-seq datasets from Cistrome database [21], which involve 1354 TFs and 642 cells/tissues (**Supplementary Table S4**). eRNA-RBP interactions are obtained based on 518 CLIP-seq datasets from POSTAR3 database [25], which involve 221 RBPs and 34 cells/tissues (**Supplementary Table S6**). TFs and RBPs with peaks located within eRNA regions are defined as potential regulators of eRNAs. E-P loops identified by 200 HiChIP experiments across 108 cell types (**Supplementary Table S7**) are collected from HiChIPdb [26]. The loops harboring anchors overlapped with query eRNAs are defined as eRNA-mediated loop.

#### 4. Subnetwork extraction

eRNA-Anno extracts hubs/modules from the overall network to obtain the tightly connected PCGs of query eRNAs. Module extraction uses SPICi [27] in the unweighted mode (default parameter).

#### 5. Functional enrichment analyses

Functional enrichment analyses, including gene ontology (GO), KEGG pathway, and MSigDB hallmark enrichment [28], are performed based on hypergeometric test using our in-house scripts (https://github.com/zhangyw0713/FunctionEnrichment).

## Results

### eRNA-ID for eRNA identification

eRNA-ID is designed for eRNA identification based on *de novo* assembled transcriptome. As shown in the input interface (http://bioinfo.szbl.ac.cn/eRNA_IDO/eRNA-ID), users need to upload a transcriptome profile in GTF format, which can be generated from RNA-seq and GRO-seq data, and define enhancer regions using our marker buffet or by uploading their BED file. eRNA-ID adopts the similar analytical workflow used in ncFANs-eLnc [12] to predict eRNAs (see **Materials and Methods**). The major advantage of eRNA-ID compared to ncFANs is the inclusion of a pre-built buffet of 8 kinds of enhancer markers (H3K27ac, H3K4me1, chromatin accessibility, RNA polymerase II binding, SEdb v2.0 super-enhancers, and three types of enhancer annotations from EnhancerAtlas v2.0 [17], FANTOM5 [18], and SCREEN [19] databases), which enables users to personalize enhancer regions of their interests. For example, users may require high-confidence enhancer regions simultaneously labeled by multiple markers or want to obtain as many enhancers as possible by merging all markers. The processing procedure of eRNA-ID is fast; a GRO-seq derived transcriptome with 3483 transcripts (SRA008244) and a total RNA-seq derived de novo transcriptome with 222,848 transcripts (GSM2824220) cost 45 and 88 seconds respectively (default parameters).

In the output interface of eRNA-ID (**Figure 2**), a table showing chromatin coordinates, enhancers, and putative targets (adjacent genes within +/- 1Mb of eRNAs) of predicted eRNAs is provided. Users can also view the information in a genome browser based on JBrowse [29]. Moreover, users can conduct functional annotation for these novel eRNAs by clicking on the “Deliver eRNA to eRNA-Anno” button.

**Figure 2.**
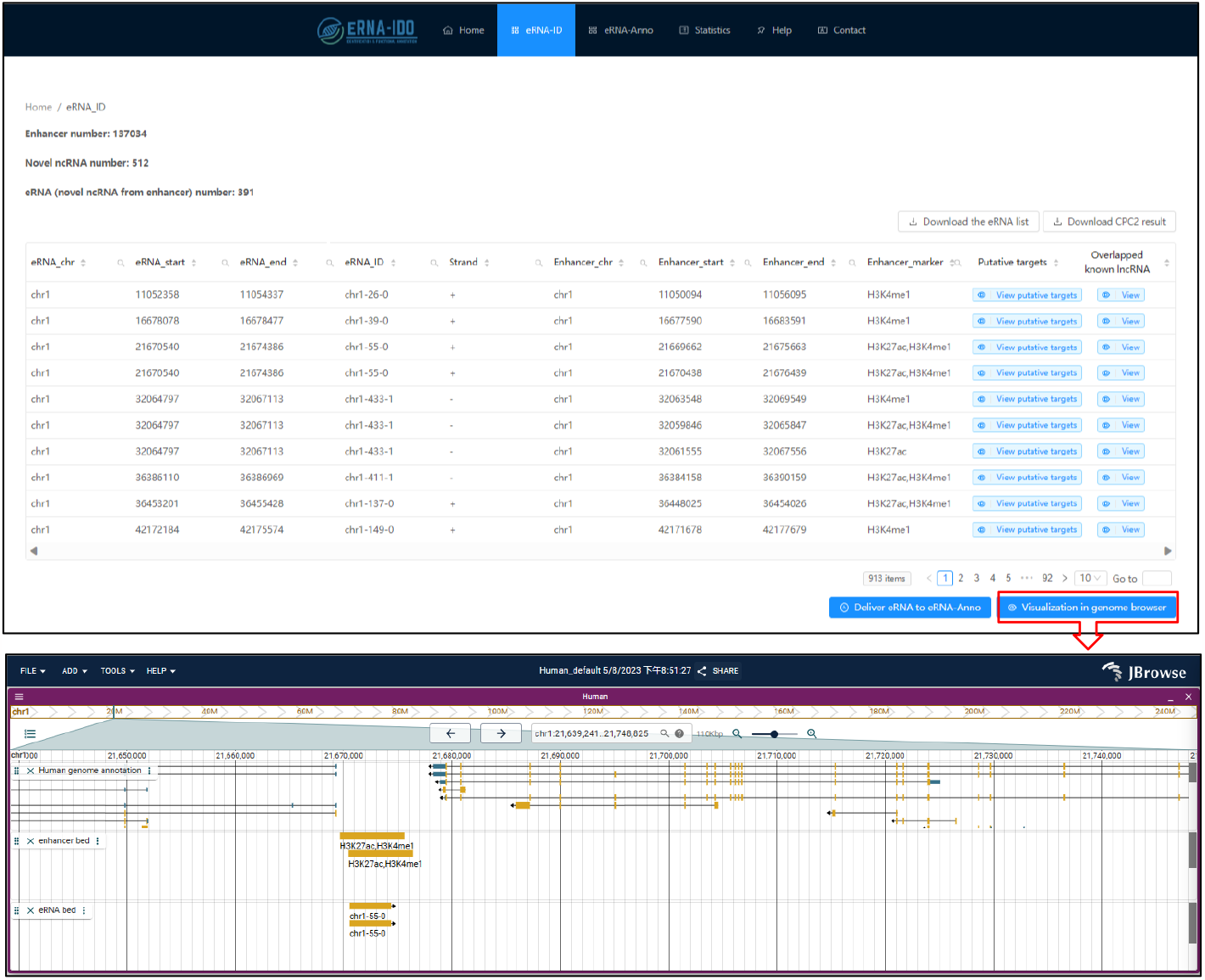
The output interface of eRNA-ID. The predicted eRNA locations, enhancer regions, markers for active enhancers, putative targets (adjacent genes), and overlapped lncRNAs are shown in a table and can be visualized in the genome browser. More details are shown in the demo: http://bioinfo.szbl.ac.cn/eRNA_IDO/retrieve/?taskid=5a9LFXS8oGCm.

### eRNA-Anno for interactome discovery and functional annotation

eRNA-Anno is designed for the network-based interactome discovery and functional annotation of eRNAs. In this module, users need to input either chromatin coordinates of novel eRNAs (**Figure 3A**) or the identifiers/locations of known eRNAs annotated in the HeRA [8] and eRic [11] database (**Figure 3B**), followed by network selection and parameter setting. eRNA-Anno first quantifies the eRNA expression levels based on RNA-seq data from TCGA and GTEx portal. As hundreds of RNA-seq samples take long processing time, we used the read coverages from BigWig files to speed up the quantification (see **Materials and Methods**). To examine the reliability of this method, we correlated the expression levels of known eRNAs based on this method with those based on canonical method and obtained from HeRA and eRic database. As expected, our method is highly correlated with canonical method using featureCounts [30] (**Figure S1A-B**) and is approximately 400 times faster (**Figure S1C**).

**Figure 3.**
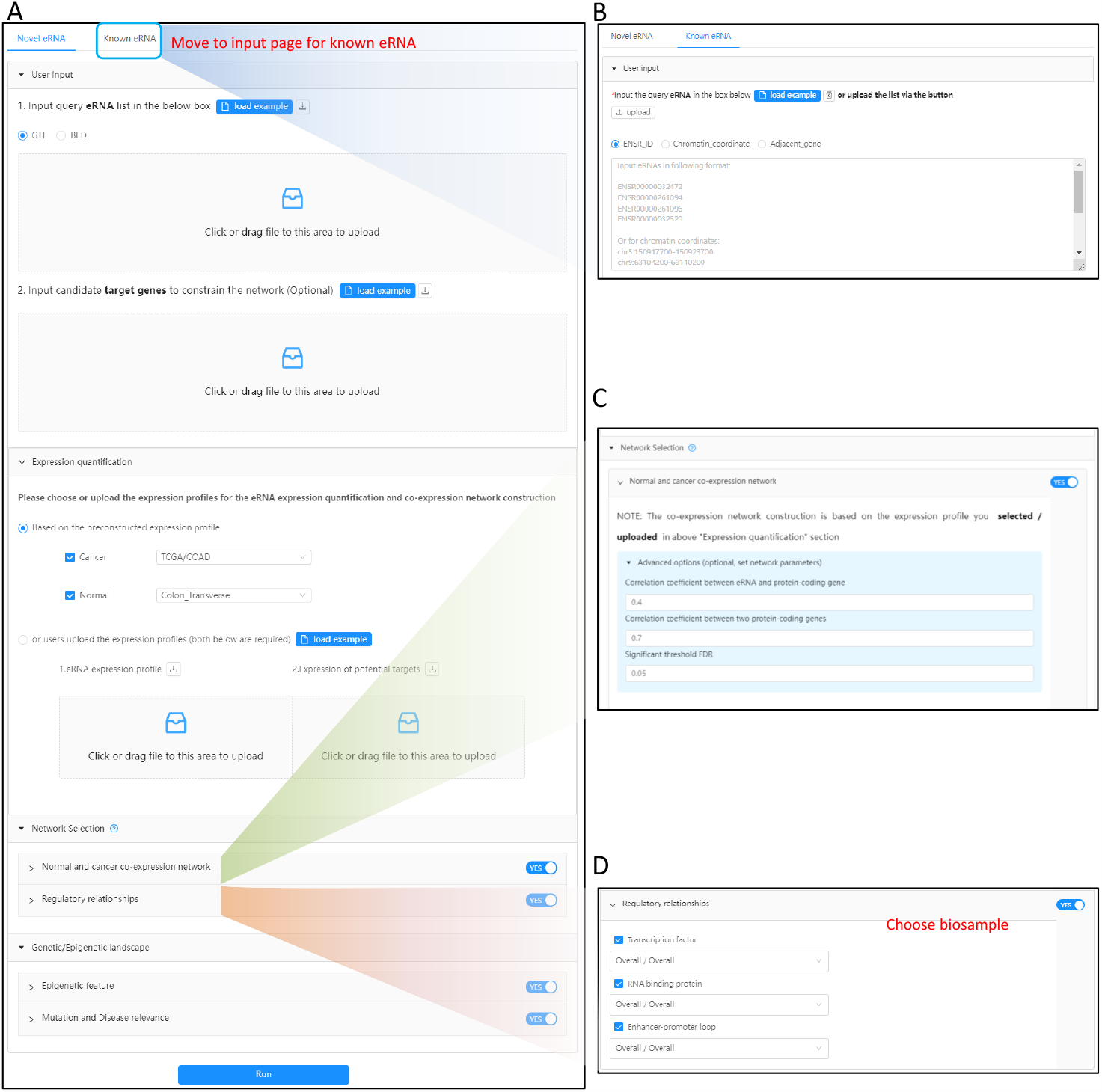
The input interface of eRNA-Anno, (A) including a potential eRNA list, optional target candidates, parameters for eRNA quantification, network selection, and genetic/epigenetic landscape. (B) Input interface for known eRNAs annotated in HeRA [8] and eRic [11]. (C) Parameters of co-expression network and (D) eRNA-centric regulatory network.

Next, eRNA-IDO annotates the functions of eRNAs through discovering their interactomes. Interactome discovery is based on the eRNA-centric networks. Networks include normal co-expression networks based on GTEx expression profiles [20], cancer co-expression networks based on TCGA expression profiles (https://portal.gdc.cancer.gov/), and eRNA-centric regulatory networks. Co-expression relationships have been widely used to annotate the functions of eRNAs [31-33]. Additionally, eRNAs were reported to exert regulatory functions through interacting with other biomolecules, including transcription factors (TFs) [34-36], RNA binding proteins (RBPs) [4, 37, 38], and target gene activated by E-P loops [39, 40]. These interactions make the regulatory network a powerful tool for eRNA functional annotation, resembling to those we used for other ncRNAs [12, 41-44]. The procedure of network construction is shown in **Materials and Methods**. Parameters include tissue/cancer type of expression profile, co-expression coefficient, significance threshold, biosamples of interaction relationships, and epigenetic landscape (**Figure 3C-D**).

Once receiving launch instruction, eRNA-Anno will initiate the analytical procedure (see **Material and Methods**) to discover the potential targets of query eRNAs from the selected networks and annotate their functions based on the hub- and module-based strategies. The whole procedure will take tens of minutes, depending on the number of the input eRNAs (**Figure. S2**). Therefore, we highly recommend users to set an email notification or record the task ID for result retrieval, when a task with a large set of eRNAs is submitted.

In the output interface, eRNA-Anno provides basic information about eRNAs (i.e., location and expression, epigenetic landscape, and disease relevance) and putative targets and functions based on different networks. In the section of “location and expression” chromatin coordinate, the expression level in normal and cancer samples, adjacent genes (<=1Mb), and overlapped super-enhancers are listed in the table (**Figure 4A**). To evaluate the activity of enhancers where eRNA is transcribed, eRNA-Anno profiles active enhancer markers (H3K27ac and H3K4me1) and chromatin accessibility of eRNA regions (**Figure 4B**). Given that mutations in eRNA regions are always related to eRNA expression and subsequent disease development [45], the clinically relevant mutations within query eRNA regions are shown (**Figure 4C**). The interactome and predicted functions of eRNAs based on the selected networks are displayed in the second part (**Figure 5**). For example, in a cancer co-expression network (**Figure 5A**), the eRNA-PCG network is visualized in a force-directed layout, and the functions of connected PCGs are provided (**Figure 5B**). Since genes with similar functions tend to be concentrically distributed, eRNA-Anno extracts hubs and modules composed of tightly connected genes from the overall network (**Figure 5D**). The function of query eRNAs can be inferred by the functions of the PCGs within the same module or hub (**Figure 5C**).

**Figure 4.**
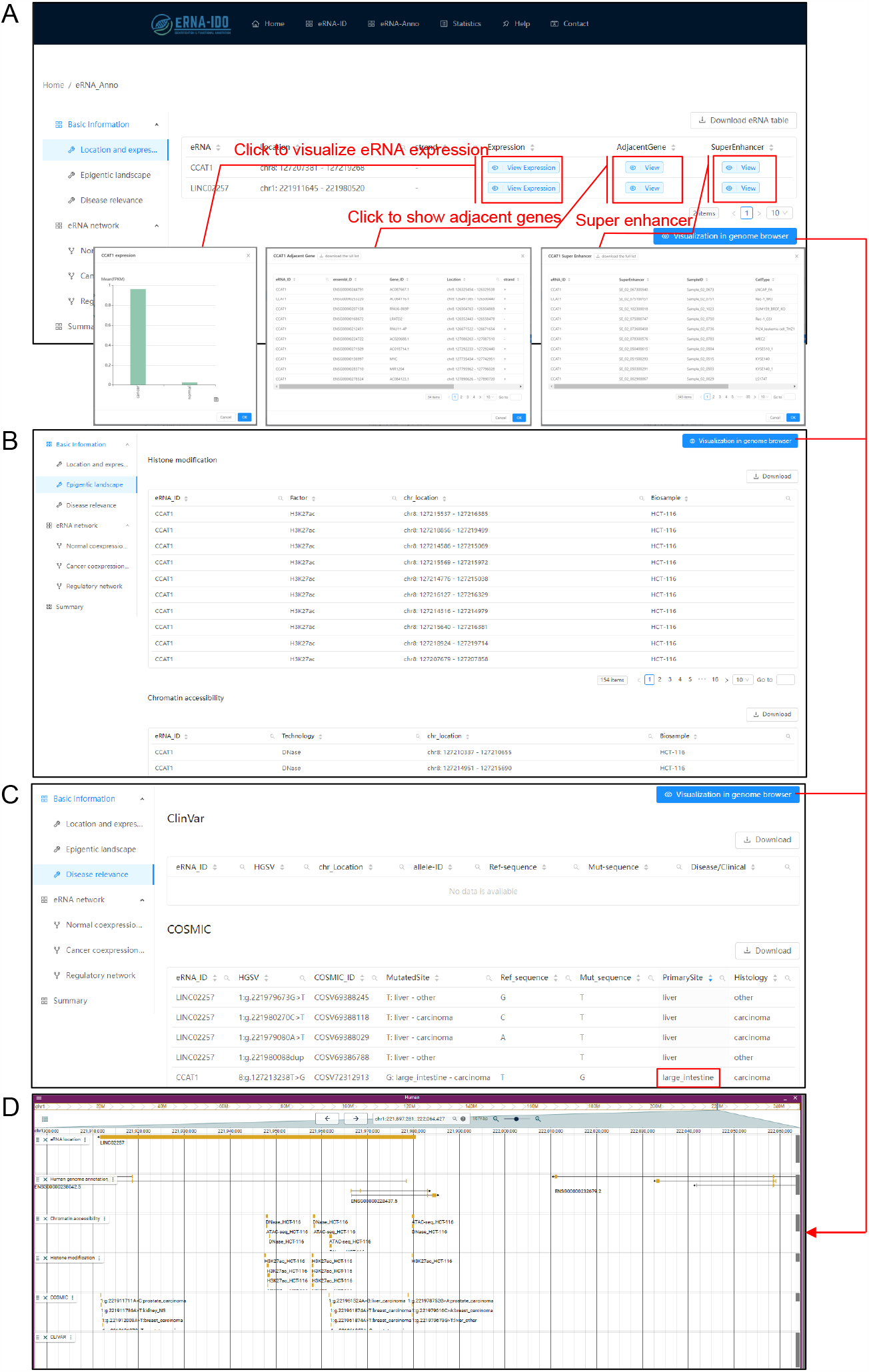
The output interface of eRNA-Anno shows the basic information of query eRNA CCAT1 and LINC02257, including (A) location and expression level, (B) epigenetic landscape, and (C) mutation-based disease relevance. (D) The genome browser can be activated by clicking on the button “Visualization in genome browser”. More details are shown in the demo: http://bioinfo.szbl.ac.cn/eRNA_IDO/retrieve/?taskid=97XPLicEAj4euYG/.

**Figure 5.**
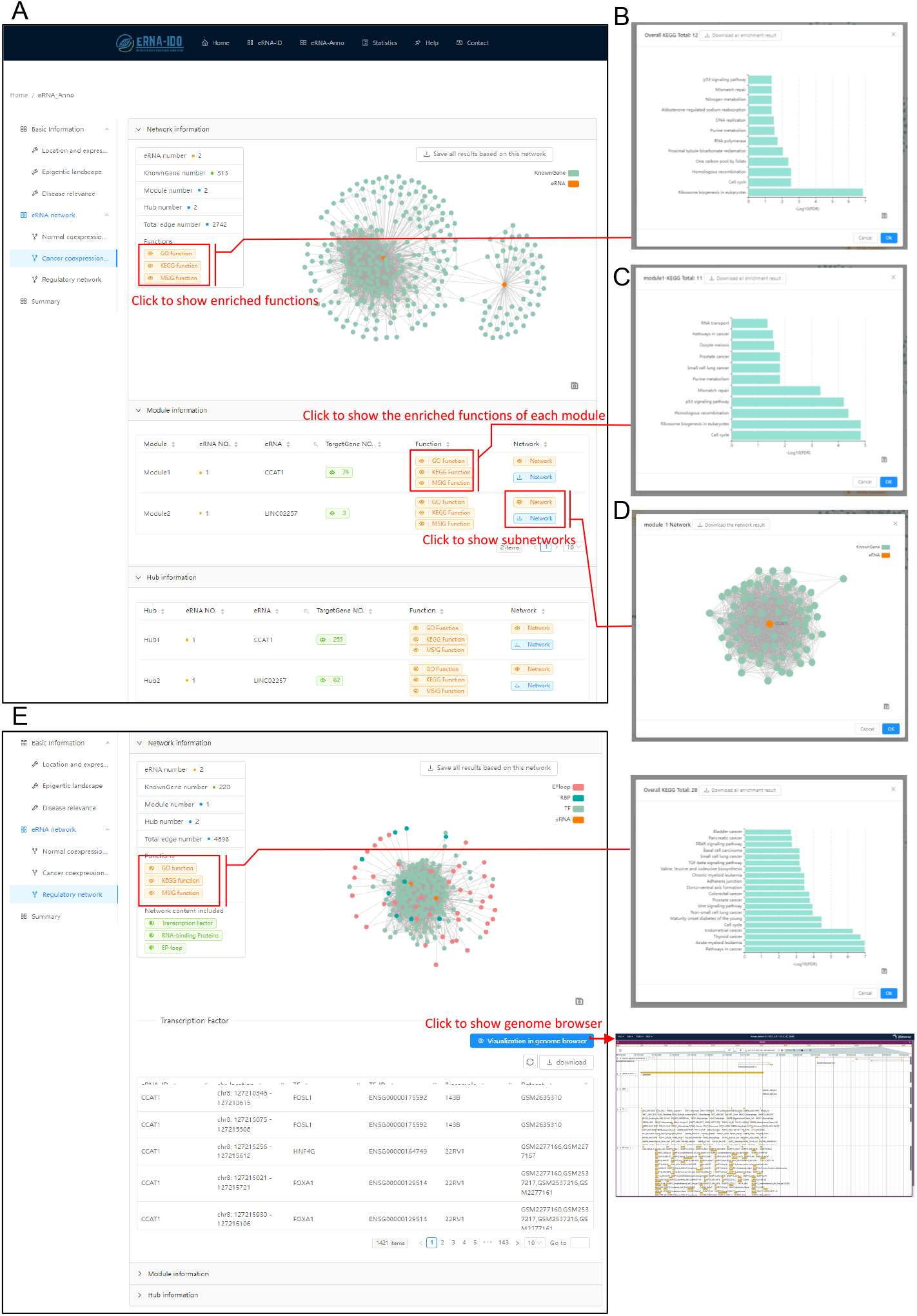
The output interface of eRNA-Anno shows the interactomes and functions of CCAT1 and LINC02257 based on (A) co-expression networks and (B) regulatory networks.

Moreover, for the eRNA-centric regulatory network (**Figure 5E)**, the relationships of eRNAs with TFs, RBPs, and E-P loops are visualized in multiple modes, including network, table, and genome browser. Similarly, the functions of eRNAs can be inferred by the related biomolecules in the overall network, modules, or hubs. After obtaining the results based on individual networks, users can combine them to get a summary (**Figure 6**).

**Figure 6.**
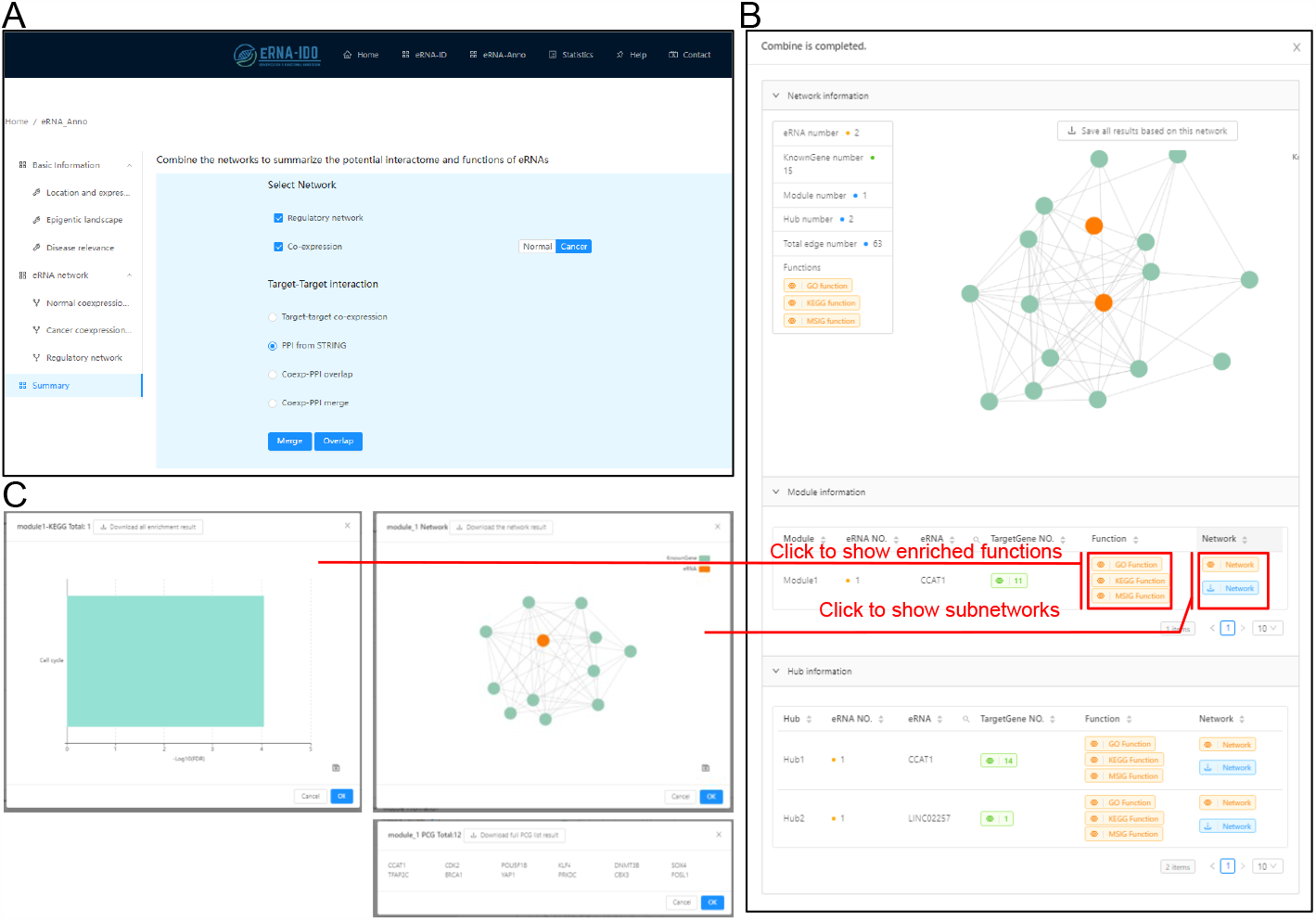
Summary of the interactome and functions of query eRNAs based on the combination of co-expression network and regulatory network. Upon (A) parameter setting, (B) a high-confidence network composed of the overlapped nodes and edges is generated for CCAT1 and LINC02257. (C) The module involved in CCAT1 indicates its interactive genes and functions in cell cycle regulation.

### A case study to showcase the usage of eRNA-Anno

As the input interface has many user-dependent options and the output interface displays interactive information, we show a case to well understand the usage and interpretation of results obtained from eRNA-Anno. CCAT1 and LINC02257, characterized as colon cancer-associated eRNAs [46, 47], were used in this study. We input them in GTF format, selected “TCGA-COAD” and “GTEx-Colon Transverse” in eRNA quantification, chose co-expression and regulatory networks, set parameters, and finally launched eRNA-Anno, as shown in **Figure 3**.

In the output interface, eRNA-Anno showed that both CCAT1 and LINC02257 exhibit higher expression levels in colorectal cancer (**Figure 4A**) and are enriched with active enhancer markers (**Figure 4B**), which is in line with the published studies [46, 47]. In addition, CCAT1 and LINC02257 regions harbor carcinoma-associated mutations (**Figure 4C**), indicating their clinical significance. To evaluate the interactome and functions of CCAT1 and LINC02257, we next looked into the co-expression network in colon adenocarcinoma. The topology of the co-expression network showed limited connections between CCAT1 and LINC02257 (**Figure 5A**), indicating their large independence in regulating target gene expression. Besides, functional enrichment analysis on the co-expressed protein-coding genes demonstrated that CCAT1 and LINC02257 are potentially enriched for translation and cell cycle pathways (**Figure 5B**). The module involved in CCAT1 precisely pinpointed the role of CCAT1 in regulating the cell cycle (**Figure 5C-D**), which conforms to previous finding [48, 49]. Moreover, the eRNA-centric regulatory network detected the interactive TFs, RBPs and the genes targeted by E-P loops, and simultaneously revealed the potential functions of CCAT1 and LINC02257 in cell cycle and cancer pathways. To intuitively visualize eRNA locations and the mutational, epigenetic and interactive landscapes, a genome browser based on JBrowse [29] was provided (**Figure 5E**). Finally, we overlapped the nodes and edges between the eRNA-centric regulatory network and cancer co-expression network. We discovered high-confidence interactions of CCAT1 in a cell cycle-related module (**Figure 6**), of which some targets such as CDK4 [50] and SOX4 [51] had been reported. This case study exemplifies the potential of eRNA-Anno, showing how it can provide comprehensive and reliable prediction on eRNA interactome and functions.

## Discussion

As a web server dedicated for eRNA, eRNA-IDO endows eRNA identification, interactome discovery and functional annotation in a convenient manner. The major advantage of eRNA-IDO include but are not limited to the below:

1. eRNA-ID provides a combination of multiple enhancer markers to realize convenient and personalized definition of enhancer regions. Compared to ncFANs-eLnc [12] with only H3K27ac marker, eRNA-ID includes 8 kinds of enhancer markers.
2. eRNA-Anno serves for any novel and known eRNAs. Considering the poor characterization of eRNAs, the applicability to novel eRNAs endows eRNA-Anno with higher flexibility and biological practicability compared to other tools requiring known identifiers such as ncFANs [12] and the databases [8-11].
3. Biological context-specific expression and interaction profiles are pre-built in eRNA-Anno. Comparing to the tools without biological specificity such as AnnoLnc2 [13], eRNA-Anno is expected to provide more precise clues for the *in vivo* investigations. Also, the pre-built profiles enable the service in a more convenient and expedite manner.
4. eRNA-IDO is the first one-stop platform for eRNA identification, interactome discovery, and functional annotation.

We also acknowledge that there remain some drawbacks and will put continuous effort to overcome them. First, our eRNA-IDO is currently designed for human data. More species will be supported in future. Second, further characteristics of eRNAs such as m^6^A modification [52] and RNA structure [53, 54] are essential for eRNA functionality but have not been investigated by eRNA-Anno. Third, current eRNA-IDO only considers normal tissue and cancer. More disease- and cell-specific expression and interaction profiles will be incorporated. Hopefully, our eRNA-IDO will benefit from user feedback and become more powerful upon our continuous updates.

## Table

**Table 1**. Data type, source, and the number of biosamples of enhancer markers

## Supplementary Figures

**Supplementary Figure 1**. Comparison of the strategies for eRNA quantification. (A-B) Distribution of Pearson correlation coefficients of eRNA levels quantified by our methods with those collected from (A) HeRA and (B) eRic database. (C) Comparison of the running time between our method and canonical featureCounts [30]. The test sample is GTEX-ZYFC-2626-SM-5NQ6S from GTEx database. The task was done on a Dell Precision T7920 workstation with single core.

**Supplementary Figure 2**. The running time of eRNA-Anno across the different eRNA numbers. X-axis and y-axis represent the input eRNA numbers and the running time, respectively.

## Supplementary Tables

**Supplementary Table 1**. ChIP-seq dataset of H3K27ac modification

**Supplementary Table 2**. ChIP-seq dataset of H3K4me1 modification

**Supplementary Table 3**. ATAC-seq/DNase-seq datasets of chromatin accessibility

**Supplementary Table 4**. ChIP-seq datasets of 1354 TFs

**Supplementary Table 5**. Normal tissue- and cancer-specific RNA-seq datasets

**Supplementary Table 6**. CLIP-seq datasets of RBPs

**Supplementary Table 7**. HiChIP datasets from HiChIPdb

## Reference

1. Lam MT, Li W, Rosenfeld MG et al. Enhancer RNAs and regulated transcriptional programs, Trends Biochem Sci 2014;39(4):170–170.

2. Han Z, Li W. Enhancer RNA: What we know and what we can achieve, Cell Prolif 2022;55(4):e13202.

3. Ounzain S, Pedrazzini T. Super-enhancer lncs to cardiovascular development and disease, Biochim Biophys Acta 2016;1863(7 Pt B):1953–1960.

4. Jiao W, Chen Y, Song H et al. HPSE enhancer RNA promotes cancer progression through driving chromatin looping and regulating hnRNPU/p300/EGR1/HPSE axis, Oncogene 2018;37(20):2728–2728.

5. Lai F, Orom UA, Cesaroni M et al. Activating RNAs associate with Mediator to enhance chromatin architecture and transcription, Nature 2013;494(7438):497–497.

6. Li W, Notani D, Ma Q et al. Functional roles of enhancer RNAs for oestrogen-dependent transcriptional activation, Nature 2013;498(7455):516–516.

7. Bose DA, Donahue G, Reinberg D et al. RNA Binding to CBP Stimulates Histone Acetylation and Transcription, Cell 2017;168(1-2):135–149 e122.

8. Zhang Z, Hong W, Ruan H et al. HeRA: an atlas of enhancer RNAs across human tissues, Nucleic Acids Res 2021;49(D1):D932–D938.

9. Chen H, Liang H. A High-Resolution Map of Human Enhancer RNA Loci Characterizes Super-enhancer Activities in Cancer, Cancer Cell 2020;38(5):701–701 e705.

10. Jin W, Jiang G, Yang Y et al. Animal-eRNAdb: a comprehensive animal enhancer RNA database, Nucleic Acids Res 2022;50(D1):D46–D53.

11. Zhang Z, Lee JH, Ruan H et al. Transcriptional landscape and clinical utility of enhancer RNAs for eRNA-targeted therapy in cancer, Nat Commun 2019;10(1):4562.

12. Zhang Y, Bu D, Huo P et al. ncFANs v2.0: an integrative platform for functional annotation of non-coding RNAs, Nucleic Acids Res 2021;49(W1):W459–W468.

13. Ke L, Yang DC, Wang Y et al. AnnoLnc2: the one-stop portal to systematically annotate novel lncRNAs for human and mouse, Nucleic Acids Res 2020;48(W1):W230–W238.

14. Frankish A, Diekhans M, Jungreis I et al. Gencode 2021, Nucleic Acids Res 2021;49(D1):D916–D923.

15. Kang YJ, Yang DC, Kong L et al. CPC2: a fast and accurate coding potential calculator based on sequence intrinsic features, Nucleic Acids Res 2017;45(W1):W12–W16.

16. Wang Y, Song C, Zhao J et al. SEdb 2.0: a comprehensive super-enhancer database of human and mouse, Nucleic Acids Res 2022.

17. Gao T, Qian J. EnhancerAtlas 2.0: an updated resource with enhancer annotation in 586 tissue/cell types across nine species, Nucleic Acids Res 2020;48(D1):D58–D64.

18. Abugessaisa I, Ramilowski JA, Lizio M et al. FANTOM enters 20th year: expansion of transcriptomic atlases and functional annotation of non-coding RNAs, Nucleic Acids Res 2021;49(D1):D892–D898.

19. Consortium EP, Moore JE, Purcaro MJ et al. Expanded encyclopaedias of DNA elements in the human and mouse genomes, Nature 2020;583(7818):699–699.

20. Consortium GT. The GTEx Consortium atlas of genetic regulatory effects across human tissues, Science 2020;369(6509):1318–1318.

21. Zheng R, Wan C, Mei S et al. Cistrome Data Browser: expanded datasets and new tools for gene regulatory analysis, Nucleic Acids Res 2019;47(D1):D729–D735.

22. Landrum MJ, Lee JM, Benson M et al. ClinVar: improving access to variant interpretations and supporting evidence, Nucleic Acids Res 2018;46(D1):D1062–D1067.

23. Tate JG, Bamford S, Jubb HC et al. COSMIC: the Catalogue Of Somatic Mutations In Cancer, Nucleic Acids Res 2019;47(D1):D941–D947.

24. Chen W, Li J, Huang S et al. GCEN: An Easy-to-Use Toolkit for Gene Co-Expression Network Analysis and lncRNAs Annotation, Curr Issues Mol Biol 2022;44(4):1479–1479.

25. Zhao W, Zhang S, Zhu Y et al. POSTAR3: an updated platform for exploring post-transcriptional regulation coordinated by RNA-binding proteins, Nucleic Acids Res 2022;50(D1):D287–D294.

26. Wang S, Gao S, Zeng Y et al. N6-Methyladenosine Reader YTHDF1 Promotes ARHGEF2 Translation and RhoA Signaling in Colorectal Cancer, Gastroenterology 2022;162(4):1183–1183.

27. Jiang P, Singh M. SPICi: a fast clustering algorithm for large biological networks, Bioinformatics 2010;26(8):1105–1105.

28. Liberzon A, Birger C, Thorvaldsdottir H et al. The Molecular Signatures Database (MSigDB) hallmark gene set collection, Cell Syst 2015;1(6):417–417.

29. Diesh C, Stevens GJ, Xie P et al. JBrowse 2: A modular genome browser with views of synteny and structural variation, bioRxiv 2022:2022.2007.2028.501447.

30. Liao Y, Smyth GK, Shi W. featureCounts: an efficient general purpose program for assigning sequence reads to genomic features, Bioinformatics 2014;30(7):923–923.

31. Cai H, Liang J, Jiang Y et al. Integrative Analysis of N6-Methyladenosine-Related Enhancer RNAs Identifies Distinct Prognosis and Tumor Immune Micro-Environment Patterns in Head and Neck Squamous Cell Carcinoma, Cancers (Basel) 2022;14(19).

32. Chen X, Yuan J, Xue G et al. Translational control by DHX36 binding to 5’UTR G-quadruplex is essential for muscle stem-cell regenerative functions, Nat Commun 2021;12(1):5043.

33. Yao P, Lin P, Gokoolparsadh A et al. Coexpression networks identify brain region-specific enhancer RNAs in the human brain, Nat Neurosci 2015;18(8):1168–1168.

34. Allen MA, Andrysik Z, Dengler VL et al. Global analysis of p53-regulated transcription identifies its direct targets and unexpected regulatory mechanisms, Elife 2014;3(e02200.

35. Azofeifa JG, Allen MA, Hendrix JR et al. Enhancer RNA profiling predicts transcription factor activity, Genome Res 2018;28(3):334–334.

36. Franco HL, Nagari A, Malladi VS et al. Enhancer transcription reveals subtype-specific gene expression programs controlling breast cancer pathogenesis, Genome Res 2018;28(2):159–159.

37. Bai X, Li F, Zhang Z. A hypothetical model of trans-acting R-loops-mediated promoter-enhancer interactions by Alu elements, J Genet Genomics 2021;48(11):1007–1007.

38. Huang Z, Yu H, Du G et al. Enhancer RNA lnc-CES1-1 inhibits decidual cell migration by interacting with RNA-binding protein FUS and activating PPARgamma in URPL, Mol Ther Nucleic Acids 2021;24(104–112.

39. Arnold PR, Wells AD, Li XC. Diversity and Emerging Roles of Enhancer RNA in Regulation of Gene Expression and Cell Fate, Front Cell Dev Biol 2019;7(377.

40. Harrison LJ, Bose D. Enhancer RNAs step forward: new insights into enhancer function, Development 2022;149(16).

41. Zhang Y, Tao Y, Li Y et al. The regulatory network analysis of long noncoding RNAs in human colorectal cancer, Funct Integr Genomics 2018;18(3):261–261.

42. Zhang Y, Tao Y, Liao Q. Long noncoding RNA: a crosslink in biological regulatory network, Brief Bioinform 2018;19(5):930–930.

43. Chen L, Zhang W, Li DY et al. Regulatory network analysis of LINC00472, a long noncoding RNA downregulated by DNA hypermethylation in colorectal cancer, Clin Genet 2018;93(6):1189–1189.

44. Luo C, Tao Y, Zhang Y et al. Regulatory network analysis of high expressed long non-coding RNA LINC00941 in gastric cancer, Gene 2018;662(103–109.

45. Hu X, Wu L, Yao Y et al. The integrated landscape of eRNA in gastric cancer reveals distinct immune subtypes with prognostic and therapeutic relevance, iScience 2022;25(10):105075.

46. McCleland ML, Mesh K, Lorenzana E et al. CCAT1 is an enhancer-templated RNA that predicts BET sensitivity in colorectal cancer, J Clin Invest 2016;126(2):639–639.

47. Xiao J, Liu Y, Yi J et al. LINC02257, an Enhancer RNA of Prognostic Value in Colon Adenocarcinoma, Correlates With Multi-Omics Immunotherapy-Related Analysis in 33 Cancers, Front Mol Biosci 2021;8(646786.

48. Liu Z, Chen Q, Hann SS. The functions and oncogenic roles of CCAT1 in human cancer, Biomed Pharmacother 2019;115(108943.

49. Li B, Zheng L, Ye J et al. CREB1 contributes colorectal cancer cell plasticity by regulating lncRNA CCAT1 and NF-kappaB pathways, Sci China Life Sci 2022;65(8):1481–1481.

50. Li JL, Li R, Gao Y et al. LncRNA CCAT1 promotes the progression of preeclampsia by regulating CDK4, Eur Rev Med Pharmacol Sci 2018;22(5):1216–1216.

51. Hu B, Zhang H, Wang Z et al. LncRNA CCAT1/miR-130a-3p axis increases cisplatin resistance in non-small-cell lung cancer cell line by targeting SOX4, Cancer Biol Ther 2017;18(12):974–974.

52. Lee JH, Wang R, Xiong F et al. Enhancer RNA m6A methylation facilitates transcriptional condensate formation and gene activation, Mol Cell 2021;81(16):3368–3368 e3369.

53. Cheng JH, Pan DZ, Tsai ZT et al. Genome-wide analysis of enhancer RNA in gene regulation across 12 mouse tissues, Sci Rep 2015;5(12648.

54. Ren C, Liu F, Ouyang Z et al. Functional annotation of structural ncRNAs within enhancer RNAs in the human genome: implications for human disease, Sci Rep 2017;7(1):15518.

