## Supplementary figures and images for "eRNA-IDO: a one-stop platform for identification, interactome discovery and functional annotation of enhancer RNAs"

### Fig.S1

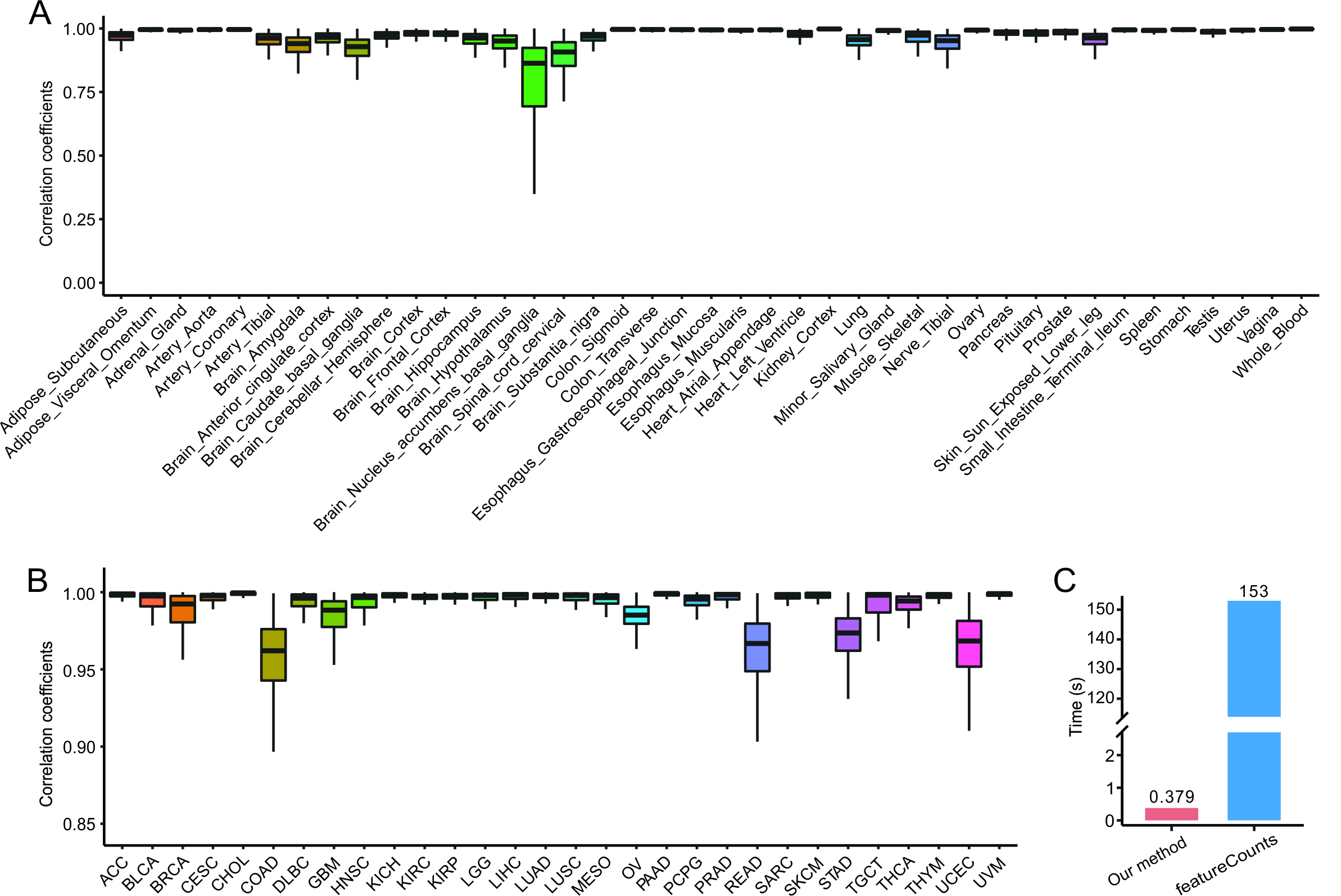

### Fig.S2

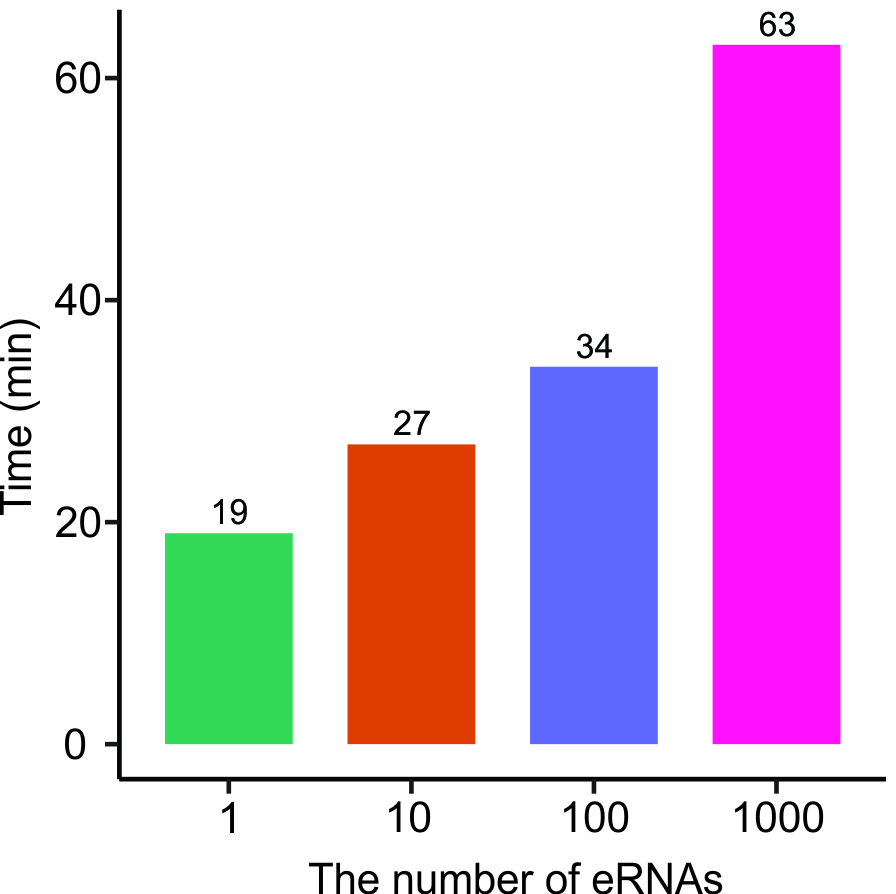
